# Apicortin defines the Plasmodium apical conoid body during transmission but is dispensable for the parasite life cycle

**DOI:** 10.1101/2025.10.01.678762

**Authors:** Mohammad Zeeshan, Akancha Mishra, Sarah L Pashley, Robert Markus, Declan Brady, Anthony A. Holder, Carolyn Moores, Rita Tewari

## Abstract

Apicomplexan parasites such as *Plasmodium* spp. and *Toxoplasma gondii* possess unique tubulin-based structures, including subpellicular microtubules and apical polar rings, which are essential for parasite motility, host cell invasion, and replication. How the stability of these structures is maintained is poorly understood, but it may involve Apicortin, a microtubule-associated protein, so-far found only in apicomplexans and the placozoan *Trichoplax adhaerens*. Apicortin contains a doublecortin (DC) domain and a partial tubulin <u>p</u>olymerisation-<u>p</u>romoting <u>p</u>rotein (TPPP) domain, both implicated in microtubule binding and stabilization. In this study, we investigated the location and function of Apicortin in *Plasmodium berghei.* Live cell imaging of a transgenic parasite line expressing GFP-tagged Apicortin showed that it was present at the apical end of invasive parasites only during development of transmission stages within the mosquito vector. High-resolution imaging using super-resolution and expansion microscopy, revealed that Apicortin forms a distinct ring-like structure in the apical complex region at the apical end of ookinetes and sporozoites. However, deletion of the Apicortin gene had no effect on parasite development and transmission through the mosquito, indicating that this protein is not essential. This suggests that there may be redundancy or compensatory functions in the mechanisms that stabilise the apical complex.

## Introduction

Plasmodium, the causative agent of malaria, is the most dangerous vector-borne protozoan parasite, responsible for an estimated 597,000 deaths in 2023 (WHO, 2024). It belongs to the phylum Apicomplexa, a group of obligate intracellular parasites, and is characterized by invasive stages - the merozoite, ookinete, and sporozoite – that are highly polarised cells. A defining feature of these cells is the presence of an anterior apical complex, which has a crucial role in parasite motility and invasion of host cells and tissues. Unlike many eukaryotic cells that use cilia or flagella for motility, apicomplexan parasites use a unique actomyosin motor known as the glideosome. This motor is located within the pellicle, a membranous structure comprised of the plasma membrane and the inner membrane complex, and drives the gliding motility essential for host cell invasion (Frenal et al., 2017).

The apical complex has several distinct structures, including an apical cap with polar rings, a subpellicular microtubule-organizing centre (MTOC), and secretory organelles including micronemes, rhoptries, and dense granules, all located on the cytosolic side of the apical complex. Another important component is the conoid — a specialized, cone-shaped structure. In some apicomplexan species, such as *Toxoplasma gondii* and *Eimeria* spp., the conoid is well defined and functional, but in other species like *Plasmodium* spp. there are only remnants of the conoid, probably reflecting a divergent adaptation of this structure (Dos Santos Pacheco et al., 2020; Koreny et al., 2021).

The apical conoid is found predominantly in apicomplexans, particularly within the coccidia, and has structural similarity to the pseudoconoid of related alveolate lineages such as *Chromera* and *Perkinsus*. In these organisms, the pseudoconoid is closely associated with the flagellar root apparatus, suggesting a common evolutionary relationship between apical complex components and flagellar structures that may have been retained or repurposed in different lineages (Portman and Slapeta, 2014).

Our recent investigation of the conoid complex revealed that many of the apical complex proteins are conserved across apicomplexan parasites (Koreny et al., 2021). Among these proteins is Apicortin (also referred to as doublecortin), which contains domains characteristic of both tubulin polymerisation-promoting proteins (TPPP) and the doublecortin (DC) family (Orosz, 2016; 2021). Although both the TPPP and DC domains are widely distributed in metazoan proteins, the presence of both domains within a single protein is an almost exclusive feature of Apicortin. This distinctive domain architecture likely reflects a specialised adaptation of the parasite’s cytoskeleton to the requirements of a highly polarised invasive cell (Orosz, 2009; 2016).

Elegant studies have described the molecular architecture of Apicortin and its role in *T. gondii* host cell invasion (Leung et al., 2020; Nagayasu et al., 2017). These investigations demonstrated that the absence of Apicortin severely impairs both host cell invasion and parasite replication (Nagayasu *et al*., 2017). Structural analyses determined the three-dimensional structure of the DC domain, and heterologously expressed Apicortin promoted the formation of microtubule (MT) bundles and short, strongly curved, arc-like structures (Leung *et al*., 2020).

*Plasmodium* species lack a defined conoid structure, but they retain a functional apical complex and encode an Apicortin homologue. However, the subcellular location and function of *Plasmodium* Apicortin are largely unknown. A recent study reported Apicortin expression in *Plasmodium falciparum* asexual blood stages, where it interacts with both α- and β-tubulins, and its repression leads to impaired host cell invasion and parasite growth (Chakrabarti et al., 2021). However, its location and function in the stages of the life cycle in the mosquito remain uncharacterised.

Here, we report on the expression dynamics and function of Apicortin in *Plasmodium berghei,* a rodent malaria model which enables the investigation of all invasive stages in the life cycle — merozoites, ookinetes, and sporozoites. We demonstrate that Apicortin is expressed in the invasive stages present only within the mosquito vector. Here, Apicortin is associated with the apical complex, forming a distinct ring-like structure. Despite this specific location of the protein, deletion of the Apicortin gene had no effect on parasite development and progression through the life cycle.

## Results

### Apicomplexan parasites have Apicortins with conserved TPPP and DC domains

We analysed the structure of Apicortin in *Plasmodium* and other apicomplexans using a range of bioinformatic tools including InterPro, Clustal Omega and AlphaFold. *P. berghei* Apicortin has a TPPP and a DC domain **(Fig. 1A)** and there is moderate sequence homology between *P. berghei*, *P. falciparum* and *T. gondii* Apicortins **(Fig. 1B).** Apicortins display a distinctive domain arrangement and overall limited sequence identity **(Fig. 1C),** including within the structurally well-defined DC domain **(Fig. 1D).**

**Fig 1.**
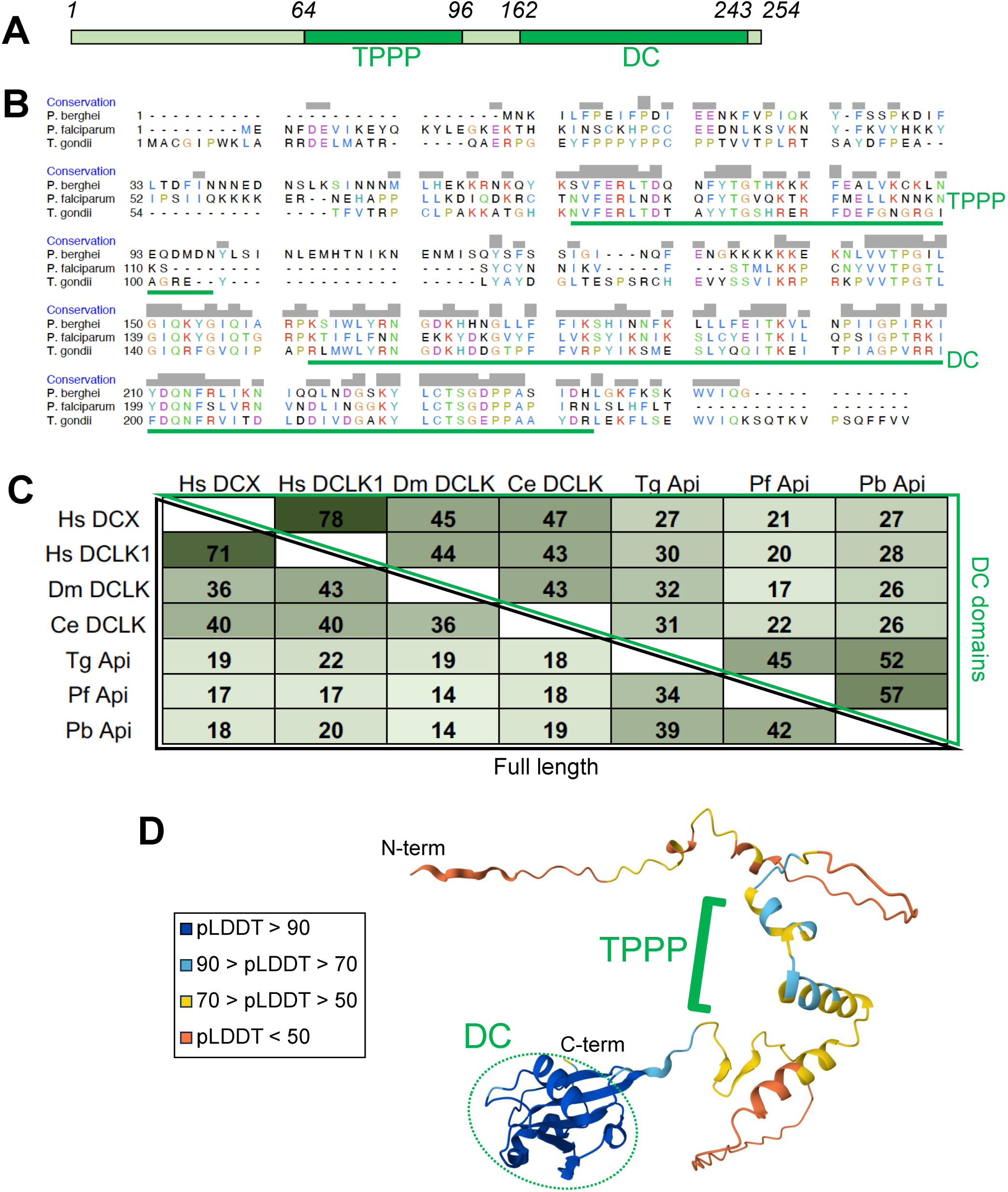
Sequence and structural analysis of Apicortins. (**A**) Schematic of *P. berghei* Apicortin (PBANKA_1232600; Uniprot: A0A509AMB2) with residue numbers indicating Uniprot domain boundaries. TPPP: Pfam PF05517; DC: Pfam PF03607. (**B**) Clustal Omega protein sequence alignment of *P. berghei, P. falciparum* (Uniprot: C0H4E4) and *T. gondii* (Uniprot: A0A7J6K3U3) Apicortin with putative microtubule-binding domains indicated. (**C**) Sequence conservation of selected DC-containing proteins with full length sequence similarity depicted on the left of the matrix (black outline) and DC domain similarity on the right (green outline). Hs DCX, *H. sapiens* doublecortin, Uniprot: O43602; Hs DCLK1, *H. sapiens* doublecortin-like kinase 1, Uniprot: O15075; Dm DCLK, *D. melanogaster* doublecortin-like kinase, Uniprot: Q7PL17; Ce DCLK *C. elegans* doublecortin-like kinase, Uniprot: Q95QC4. (**D**) AlphaFold predicted structure of *P. berghei* Apicortin, coloured according to the per-residue confidence metric, pLDDT from 1-100, and with TPPP and DC domains and N- and C-termini indicated.

### *Pb*Apicortin is located at the apical end of invasive parasites in the mosquito vector

To investigate the expression and location of *Plasmodium* Apicortin, we generated a transgenic *P. berghei* line expressing Apicortin with a C-terminal GFP tag. An in-frame *gfp* coding sequence was inserted at the 3’ end of the endogenous *apicortin* locus using single crossover homologous recombination (**Fig S1A**). Successful insertion was confirmed by diagnostic PCR (**Fig S1B**).

Both asexual blood stages in the mammalian host and sexual stages in the mosquito of this PbApicortin-GFP transgenic parasite line were examined for the expression of Apicortin, using live-cell imaging. Expression was undetectable during asexual blood-stage replication, in both male and female gametocytes, and in flagellated male gametes. PbApicortin-GFP was first detected at the zygote stage, approximately two hours after fertilisation, located at the cell periphery as a single focal point **(Fig 2A).** To follow the location of PbApicortin-GFP during zygote differentiation, we examined the different stages of zygote development (stages I–V) to the fully motile and invasive ookinete (stage VI) (**Fig 2A**). PbApicortin-GFP was detected at a single focal point during stage 1 on the cellular protrusion defining the apical polarity that develops as the zygote starts to transform into a zoite. As the zoite elongated through stages II to V, PbApicortin-GFP remained located at the polar end of the protrusion and eventually at the apical end of the mature ookinete (stage VI) (**Fig 2A**).

**Fig. 2.**
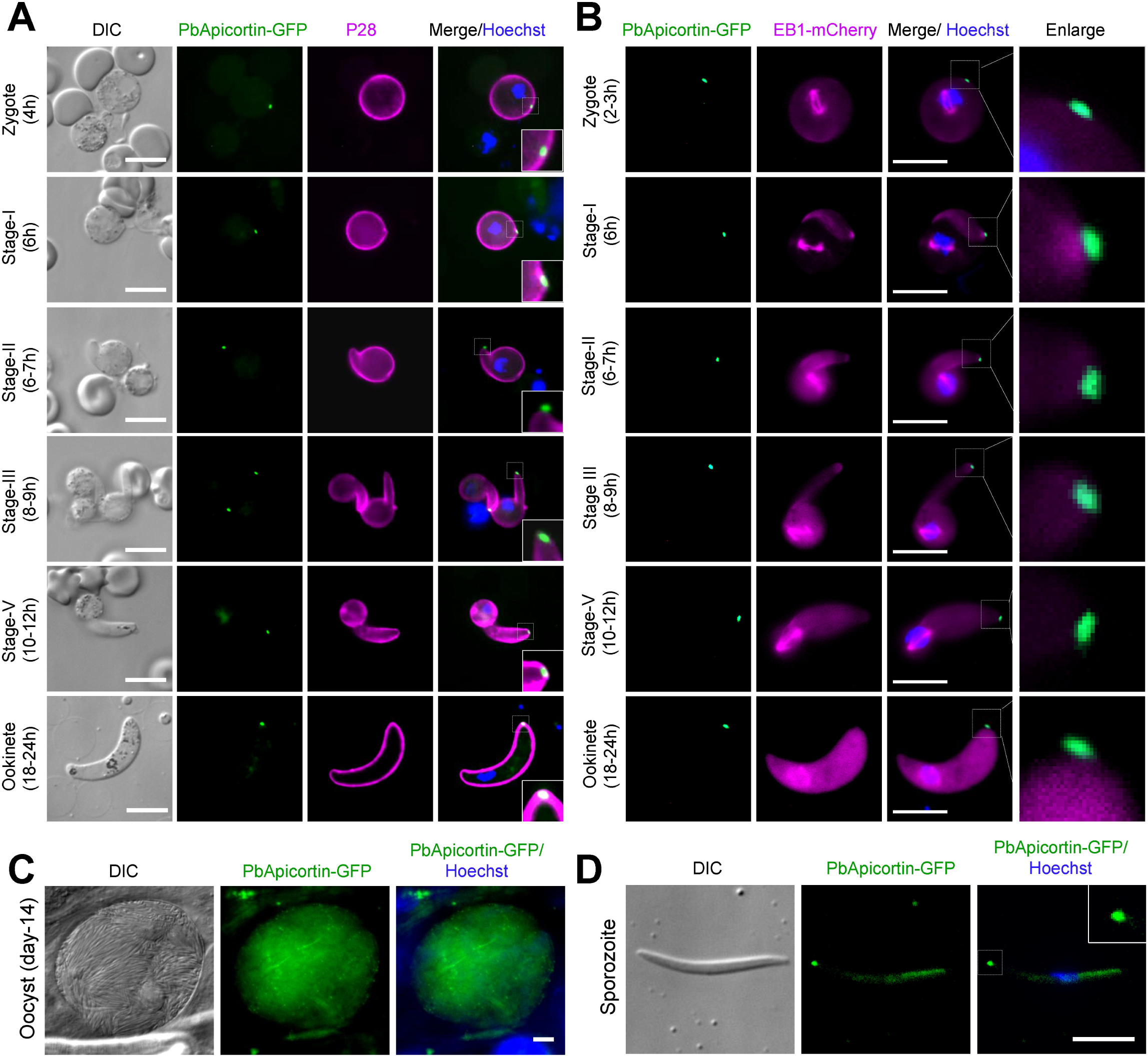
*P. berghei* Apicortin has an apical location during zygote transformation to ookinete and during sporogony. **(A)** Live cell imaging showing *Pb*Apicortin-GFP location during zygote to ookinete transformation. A cy3-conjugated antibody, 13.1, which binds to the P28 protein on the surface of activated female gametes, zygotes and ookinetes was used to mark these stages (magenta). Panels: DIC (differential interference contrast), PbApicortin-GFP (green, GFP), Merged: Hoechst (blue, DNA), PbApicortin-GFP (green, GFP) and P28 (magenta). Scale bar=5 μm. Insets show the zoom of PbApicortin-GFP signal. **(B)** Live cell imaging showing the location of PbApicortin-GFP (green) with respect to EB1-mCherry (magenta) during zygote to ookinete transformation. Scale bar=5 μm. **(C)** Live cell imaging of PbApicortin-GFP in oocysts in mosquito guts at 14-days post-infection Panels: DIC (differential interference contrast), Apicortin-GFP (green, GFP), Merged: Hoechst (blue, DNA) and PbApicortin-GFP (green, GFP). Scale bar = 5 µm. **(D)** Live cell imaging of PbApicortin-GFP in sporozoites. Panels: DIC (differential interference contrast), PbApicortin-GFP (green, GFP), Merged: Hoechst (blue, DNA) and PbApicortin-GFP (green, GFP). Scale bar = 5 µm

To compare the location of PbApicortin-GFP relative to that of *P. berghei* End-binding microtubule protein 1 (PbEB1), a protein known to be located at the apical end of the ookinete, we genetically crossed PbApicortin-GFP (green) and PbEB1-mCherry (red) parasite lines to obtain parasites expressing both fluorescent markers. *Pb*EB1 is located at the apical end of the cell during ookinete development (Zeeshan et al., 2023) as well as showing lateral attachment to spindle MTs in the nucleus (Yang et al., 2023; Zeeshan *et al*., 2023). Live cell imaging of these parasites showed that the location of Apicortin (green) is more apical than that of PbEB1 (magenta) during ookinete development (Stages I-IV) **(Fig 2B).** In later more mature ookinete stages the apical PbEB1-mCherry signal becomes progressively diffuse and finally disappears, but the apical PbApicortin-GFP signal remains until the end of ookinete development **(Fig 2B).**

The mature ookinete transverses the mosquito gut wall to form an oocyst on the outer surface of gut wall, where multiple nuclear divisions produce thousands of sporozoites (Guttery et al., 2022). Therefore, we investigated the expression of Apicortin during this oocyst development. We observed a diffused location of PbApicortin-GFP in oocysts with some distinct foci (**Fig 2C**). The sporozoites that emerged from these oocysts at day-14 following mosquito infection, had a strong apical focal point of PbApicortin-GFP expression, together with a diffused distribution within the sporozoite (**Fig 2D**). A similar patten of PbApicortin-GFP distribution was seen in salivary gland sporozoites.

### Ultrastructural analysis of *Pb*Apicortin shows a ring-like structure at the apical end of ookinetes and sporozoites

To further examine the location of Apicortin, structured illumination microscopy (3D-SIM) was performed on fixed developing ookinetes (stage II) and fully mature ookinetes (stage VI). We found that the focal point of PbApicortin-GFP observed in wide field microscopy is a ring-like structure at the apical end of the ookinete protrusion **(Fig 3A)**, suggesting that Apicortin is part of a conoid remnant, consistent with our previous prediction (Koreny *et al*., 2021).

**Fig. 3.**
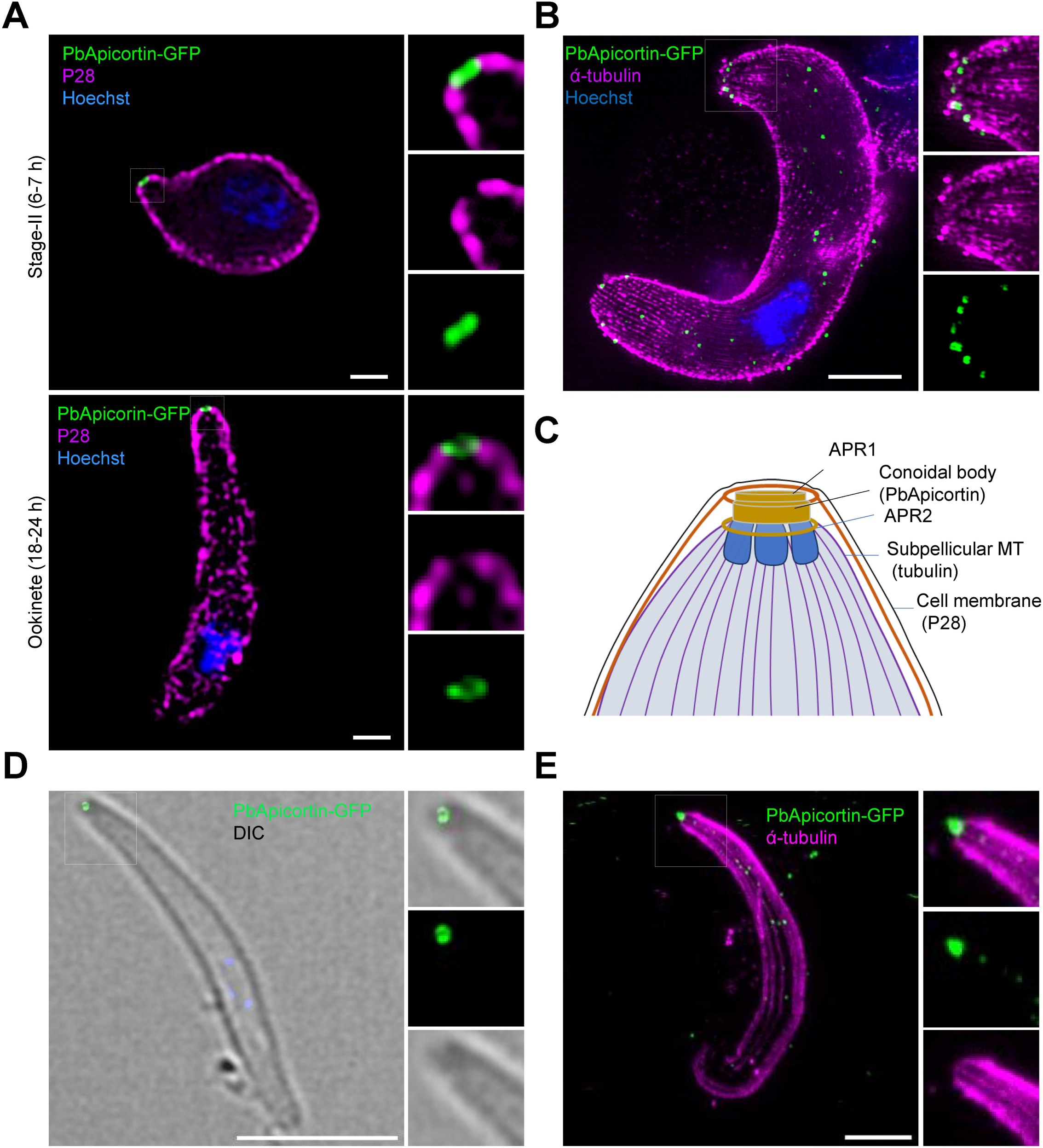
Ultrastructural analysis reveals a PbApicortin ring-like structure associated with subpellicular microtubules. **(A)** 3D-SIM images showing PbApicortin-GFP location during ookinete differentiation (stage II and VI). Hoechst (blue, DNA), PbApicortin-GFP (green, GFP) and P28 (magenta). Scale bar = 1 µm. **(B)** U-ExM image showing ring-like structure of PbApicortin-GFP (green) and subpellicular MTs (magenta) stained with alpha-tubulin antibody. Scale bar=5 μm. **(C)** Schematic of the proposed apical region of an ookinete showing the location of Apicortin. **(D)** 3D-SIM images showing PbApicortin-GFP location in sporozoites. Hoechst (blue, DNA), PbApicortin-GFP (green, GFP). Scale bar = 5 µm. **(E)** U-ExM image of sporozoite showing the ring-like structure of PbApicortin (green) and subpellicular MTs (magenta) stained with alpha-tubulin antibody. Scale bar=5 μm.

To define in more detail the ultrastructure, we analysed the location of PbApicortin-GFP relative to subpellicular MTs using ultrastructure expansion microscopy (U-ExM) analysis of ookinetes. Subpellicular MTs are organised at apical ring 2 (APR2) (Koreny *et al*., 2021) in Plasmodium and play a crucial role in determining the shape, size, and polarity of the parasite cell. U-ExM images revealed Apicortin as a ring-like structure at the apical end of the ookinete **(Fig 3B),** consistent with the 3D-SIM observation (**Fig 3A)**. Subpellicular MTs stained with antibody to alpha-tubulin were visible appearing from the Apicortin-ring **(Fig 3B).**

Together, live-cell imaging and ultrastructural analysis of ookinetes indicate that Apicortin forms a ring-like structure at the apical end of the cell, distal to EB1 and connected to the subpellicular MTs, as summarised in the schematic **(Fig. 3C**). Previously we had proposed that Apicortin is part of the conoid body (Koreny *et al*., 2021) and our current finding supports this suggestion. The schematic (**Fig 3C**) shows the arrangement of the apical complex and its different structures, with the possible location of Apicortin highlighted, based on a previous prediction and current findings with ookinetes.

To examine in more detail the location of Apicortin in sporozoites within the mosquito, 3D-SIM and expansion microscopy were used. A similar location was observed by both 3D-SIM **(Fig 3D)** and U-ExM **(Fig 3E)** in sporozoites, consistent with the proposed location of Apicortin at the apical ring of these invasive stages.

### Apicortin is not necessary for parasite transmission by mosquitoes

The function of Apicortin during the *Plasmodium* life cycle was examined by deleting its gene from the *P. berghei* genome. This was achieved using a double crossover homologous recombination strategy, using a parasite line that constitutively expresses GFP throughout the parasite life cycle (**Fig. S1C**) (Janse et al., 2006). The successful integration of the targeting construct at the *apicortin* locus was confirmed by integration PCR (**Fig. S1D**). Successful creation of this transgenic parasite (Δ*PbApicortin)* indicated that the gene is not essential for the asexual blood stages. Further phenotypic analysis of the Δ*apicortin* parasite compared with the parental parasite (WT-GFP) was carried out at other stages of the life cycle.

Δ*PbApicortin* and WT-GFP parasites produced a comparable number of gametocytes in mice, so we next analysed male and female gametocyte differentiation. Male gamete development in *Plasmodium* is a rapid process with three rounds of genome duplication and axoneme formation resulting in eight motile flagellated gametes, in a process named exflagellation (Sinden et al., 1976). There was no defect in male gamete exflagellation and female gamete formation in Δ*PbApicortin* parasite comparing to WT-GFP (**Fig. 4A**). Zygote formation and ookinete differentiation were also similar in Δ*PbApicortin* and WT-GFP parasites (**Fig. 4B**). The shape and motility of ookinetes were also not affected by the Apicortin gene deletion (**Fig 4C, D, Videos SV1 and SV2**). Detailed structural analysis by U-ExM revealed no morphological differences in apical polarity and MT organisation in Δ*PbApicortin* ookinetes **(Fig 4E).**

**Fig. 4.**
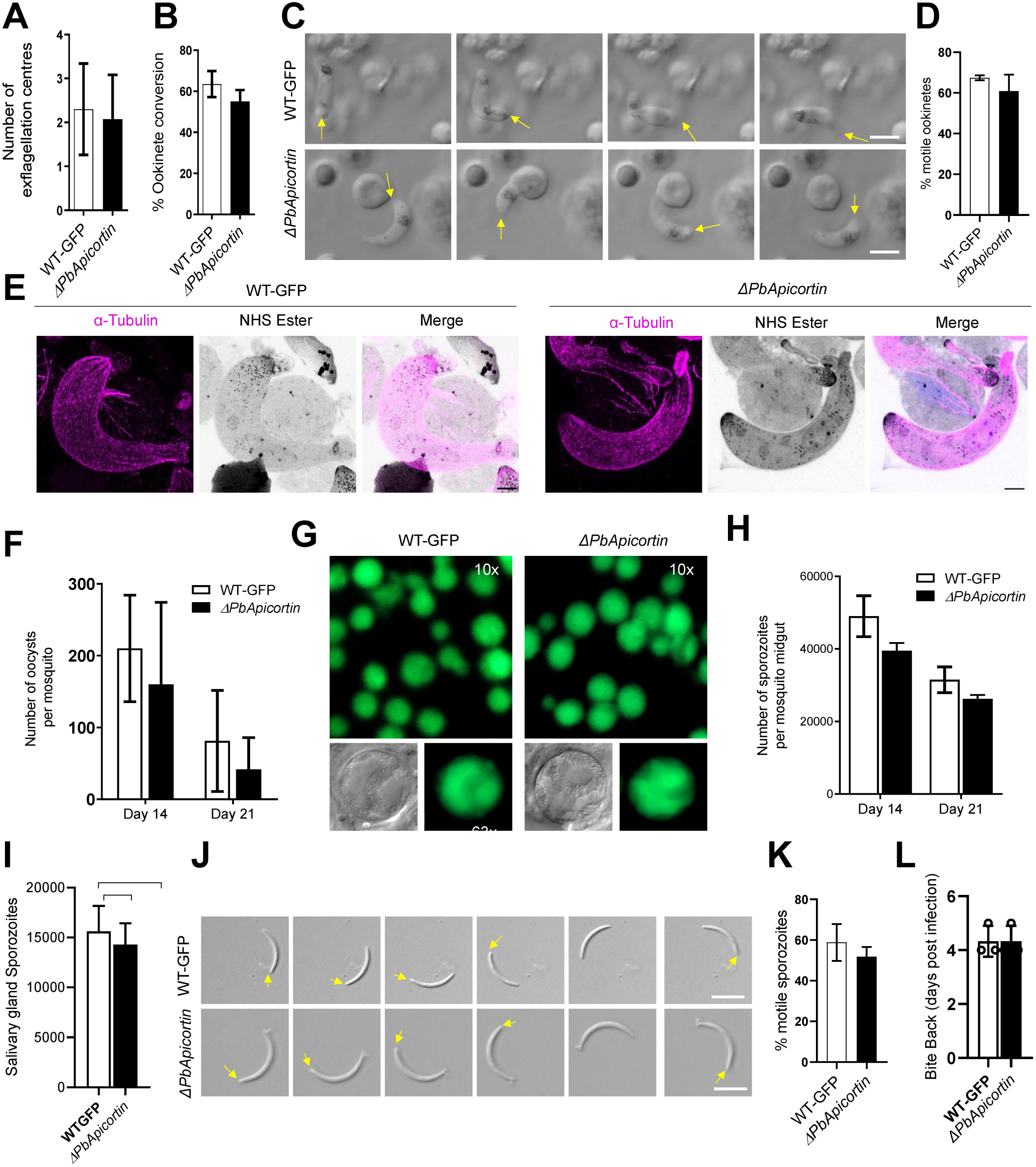
PbApicortin is not necessary for parasite transmission. **(A)** Male gametogony (exflagellation) of Δ*PbApicortin* (black bar) and WT-GFP (white bar) lines measured as the number of exflagellation centres per field. Mean ± SD; n=3 independent experiments. **(B)** Ookinete conversion as a percentage for Δ*PbApicortin* (black bar) and WT-GFP (white bar) parasites. Ookinetes were identified using 13.1 antibody as a surface marker and defined as those cells that differentiated successfully into elongated ‘banana shaped’ ookinetes. Mean ± SD; n=3 independent experiments. **(C)** Differential interference contrast (DIC) time-lapse image sequences showing motile Δ*PbApicortin* and WT-GFP ookinetes. **(D)** Quantitative data for motile ookinetes for Δ*PbApicortin* and WT-GFP based on two independent experiments. **(E)** U-ExM images showing ookinetes labelled with anti-tubulin antibody and NHS ester. Scale bar = 5 μm. **(F)** Total number of GFP-positive oocysts per infected mosquito in Δ*PbApicortin* (black bar) and WT-GFP (white bar) parasites at 14- and 21-days post infection (dpi). Mean ± SD; n=3 independent experiments. **(G)** Mosquito mid guts at 10x and 63x magnification showing oocysts of Δ*PbApicortin* and WT-GFP lines at 14 dpi. Scale bar = 50 μm in 10x and 20 μm in 63x. **(H)** Total number of sporozoites in oocysts of Δ*PbApicortin* (black bar) and WT-GFP (white bar) parasites at 14 and 21 dpi. Mean ± SD; n=3 independent experiments. **(I)** Total number of sporozoites in salivary glands of Δ*PbApicortin* (black bar) and WT-GFP (white bar) parasites. Bar diagram shows mean ± SD; n=3 independent experiments. **(J)** Differential interference contrast (DIC) time-lapse image sequences showing motile Δ*PbApicortin* and WT-GFP sporozoites isolated from salivary glands. Arrow indicates apical end of sporozoites. Scale bar = 5 μm. **(K)** Quantitative data for motile sporozoites from salivary glands for Δ*PbApicortin* and WT-GFP lines based on two independent experiments. **(L)** Bite back experiments show successful transmission of Δ*PbApicortin* parasites (black bar) from mosquito to mice, like WT-GFP parasites (white bar). Mean ± SD; n= 3 independent experiments.

To investigate any loss of function in oocyst development and sporogony resulting from deletion of the Apicortin gene, *Anopheles stephensi* mosquitoes were fed on mice infected with Δ*PbApicortin* parasites or WT-GFP parasites as a control. The number of GFP-positive oocysts on the mosquito gut wall was counted on days 14 and 21 post-infection. There was no significant difference in the number of Δ*PbApicortin* and WT-GFP oocysts ((**Fig. 4F**), and the size of the oocysts was similar for both parasites (**Fig. 4G**). There was no significant difference in Δ*PbApicortin* sporozoite numbers in oocysts at days 14 and 21 post-infection, or in salivary glands at day 21 compared to WT-GFP parasites (**Fig. 4H, I**). The shape, size, and motility of salivary gland sporozoites were indistinguishable from those of WT-GFP parasites (**Fig. 4J, K, videos SV3 and SV4)**. When infected mosquitoes were used in bite back experiments to ascertain the infectivity of Δ*PbApicortin* sporozoites in mice, a blood stage infection was observed after 4 days with both Δ*PbApicortin* and WT-GFP sporozoites (**Fig. 4L**) indicating that Apicortin is not necessary for parasite transmission through the mosquito.

## Discussion

In this study, we provide new insights into the sequence conservation, structural features, and functional relevance of Apicortins in *Plasmodium* and other apicomplexan parasites. Through a combination of comparative sequence analysis and AlphaFold-based structural modelling, we show that Plasmodium Apicortins retain a conserved architecture featuring TPPP and DC domains, consistent with the microtubule/tubulin-associated role of Apicortin in the *T. gondii* tachyzoite apical conoid structure (Leung *et al*., 2020; Nagayasu *et al*., 2017; Sun et al., 2022). Both TPPP and DC domains are microtubule or tubulin polymer-binding modules (Orosz, 2021), and despite only moderate sequence conservation, particularly within the DC domain, the maintenance of this distinct domain combination in apicomplexans suggests a specialised role of Apicortin in the unique conoid structures of these parasites.

We demonstrated the location of Apicortin at the apical end of ookinetes and sporozoites as a ring-like structure during these invasive parasite stages in the mosquito. In ookinetes the apical location is established early during zygote development and persists through ookinete maturation. Its location is maintained in both midgut and salivary gland sporozoites. The location of Apicortin implicates it in apical complex organisation throughout parasite transmission stages within the mosquito.

A microtubule-associated role for proteins that contain TPPP domains (also referred to as p25alpha domains) has been proposed in a number of physiological contexts in humans, including a role in myelination of oligodendrocytes (Fu et al., 2019), and linked to the neurodegenerative features of both Alzheimer’s and Parkinson’s diseases (Olah et al., 2011). DC domains were first characterised in human doublecortin (DCX), a protein required for neuronal migration during brain development. Doublecortin and related doublecortin-like kinases are conserved in metazoa and are characterised by tandem DC domains (N-DC and C-DC), both of which can bind MTs and contribute to a range of microtubule-related functions (Reiner et al., 2006).

The distinct domain arrangement and modest sequence identity of Apicortins even in the structurally well-predicted DC domain, argues that these proteins be considered as unique. This distinction is further supported by the specialised fibrillar arrangement of tubulin within the apicomplexan conoid complex – referred to as conoid fibrils or the conoid canopy – which is highly distorted compared to cellular MTs (Hu et al., 2002; Sun *et al*., 2022). Strikingly, both TPPP and DC domains bind to MTs between the protofilaments from which they - and the conoid fibrils - are built (Legal et al., 2024; Moores et al., 2004). This inter-protofilament binding site differs subtly in MTs with differing protofilament numbers (Moores et al, 2004), and may be dramatically modified by specific binding proteins, including Apicortin, to form the conoid complex.

PbApicortin expression was not detected in blood stage merozoites, consistent with the absence of other conoid-associated proteins that are restricted to ookinetes and sporozoites (Wall et al., 2016; Koreny et al., 2022). This stage-specific restriction suggests that Apicortin contributes to the structural or functional properties of the apical complex that are uniquely required for parasite development within the mosquito vector, but dispensable during erythrocyte invasion. In contrast, some conoid-associated proteins are expressed across all three invasive stages: ookinetes, sporozoites and merozoites (Koreny et al., 2022), highlighting the idea that while there is a conserved apical complex scaffold, the composition may be remodelled to meet the distinct demands of different life-cycle stages. For example, merozoites invade host cells, but both ookinetes and sporozoites must invade and traverse host tissue.

Genetic deletion of *P. berghei* Apicortin had no detectable consequence for parasite differentiation, proliferation, or transmission through the mosquito vector. This may be because Apicortin function is redundant for conoid formation or function, or because conoid disruption is insufficient to prevent *P. berghei* transmission.

These findings support the idea that Apicortin is a structural component of the apical complex, contributing to the unique microtubule arrangements within the conoid-like structures of *Plasmodium* ookinetes and sporozoites. Given their MT-binding domain configuration and location, Apicortins may facilitate the assembly or stabilisation of the highly curved protofilament arrangements characteristic of apicomplexan apical complexes.

## Material and Methods

### Ethics statement

The animal work performed in this study has passed an ethical review process and was approved by the United Kingdom Home Office. Work was carried out under UK Home Office Project Licenses (PDD2D5182; PP3589958) in accordance with the UK ‘Animals (Scientific Procedures) Act 1986’. Six-to eight-week-old female CD1 outbred mice from Charles River laboratories were used for all experiments. The mice were kept in a 12h light and 12h dark (7 am till 7 pm) cycle, at a temperature between 20-24 °C and the humidity between 40-60%.

### Generation of transgenic parasites

To generate the GFP-tagged lines, a region of *Pbapicortin* downstream of the ATG start codon was amplified, ligated to p277 vector, and transfected as described previously (Guttery et al., 2012). The p277 vector contains the human *dhfr* cassette, conveying resistance to pyrimethamine. Schematic representations of the endogenous gene locus, the constructs and the recombined gene locus can be found in **Supplementary Fig. S1A.** Diagnostic PCR was used with primer 1 (IntT275) and primer 3 (ol492) to confirm integration of the GFP targeting construct **(Supplementary Fig S1B).**

To delete the gene for Apicortin, the targeting vector for *apicortin* was constructed using the pBS-DHFR plasmid, which contains polylinker sites flanking a *T. gondii dhfr/ts* expression cassette conferring resistance to pyrimethamine, as described previously (Tewari et al., 2010). The 5′ upstream sequence of *apicortin* was amplified from genomic DNA and inserted into *Apa*I and *Hin*dIII restriction sites upstream of the *dhfr/ts* cassette of pBS-DHFR. A DNA fragment amplified from the 3′ flanking region of *apicortin* was then inserted downstream of the *dhfr/ts* cassette using *Eco*RI and *Xba*I restriction sites. The linear targeting sequence was released from the plasmid using *Apa*I/*Xba*I restriction enzymes. A schematic representation of the endogenous *apicortin* locus, the construct and the recombined locus can be found in **Supplementary Fig. S1C**. A diagnostic PCR with primer 1 (IntN141_5) and primer 2 (ol248) was used to confirm integration of the targeting construct, and primer 3 (KO1) and primer 4 (KO2) were used to confirm deletion of the *apicortin* gene **(Supplementary Fig. S1D)**. A list of primers used to amplify gene sequences can be found in **Supplementary Table S1**.

*P. berghei* ANKA line 2.34 (for GFP-tagging) or ANKA line 507cl1 expressing GFP (for gene deletion) parasites were transfected by electroporation (Janse *et al*., 2006).

### Live-cell imaging

Different stages of parasite development during the asexual blood stage, the zygote to ookinete transformation, and oocyst development were analysed for PbApicortin-GFP expression and location using 10x and 63x oil immersion objectives on a Zeiss Axio Imager M2 microscope and analysed with the AxioVision 4.8.2 and Zen lite 3.8 (Zeiss) software.

### Ookinete culture and purification

Mouse blood infected with PbApicortin-GFP parasites, with approximately 10% gametocytaemia, were incubated in ookinete culture medium (RPMI1640+20% FBS+ 100 µM Xanthurenic acid) for 24 h. Under these conditions the gametocytes differentiate to gametes, fertilize and produce zygotes within 30 min. The zygotes differentiate to mature ookinetes over 24 h. Developing ookinetes (at 6 to 7 h post incubation) and mature ookinetes (at 24 h) were purified using magnetic beads coated with antibodies against P28 (a surface protein of zygotes and ookinetes). The purified ookinetes were fixed with 4% paraformaldehyde (PFA) and used for structured illumination and expansion microscopy (Zeeshan et al., 2022).

### Structured illumination microscopy (3D-SIM)

The fixed ookinetes were resuspended in RPMI 1640 containing Hoechst dye and P28 antibody labelled with Cy3. Cells were scanned with an inverted microscope using Zeiss Plan-Apochromat 63x/1.4 Oil immersion or Zeiss C-Apochromat 63×/1.2 W Korr M27 water immersion objective on a Zeiss Elyra PS.1 microscope, using the structured illumination microscopy (3D-SIM) technique. The correction collar of the objective was set to 0.17 for optimum contrast. The following settings were used in SIM mode: lasers, 405 nm: 20%, 488 nm: 16%, 561nm: 8%; exposure times 200 ms (Hoechst) 100 ms (GFP for Apicortin) and 200ms (dxRed for P28); three grid rotations, five phases. The band pass filters BP 420-480 + LP 750, BP 495-550 + LP 750 and BP 570-620 + 750 were used for the blue, green and red channels, respectively. Multiple focal planes (Z stacks) were recorded with 0.2 µm step size; later post-processing, a Z correction was done digitally on the 3D rendered images to reduce the effect of spherical aberration (reducing the elongated view in Z; a process previously tested with 0.1µm fluorescent beads, ThermoFisher T7284). Registration correction was applied based on control images of multicolour fluorescent beads. Images were processed and all focal planes were digitally merged into a single plane (Maximum intensity projection). The images recorded in multiple focal planes (Z-stack) were 3D rendered into virtual models and exported as images and movies (see supplementary material). Processing and export of images and videos were done by Zeiss Zen 2012 Black edition, Service Pack 5 and Zeiss Zen 2.1 Blue edition.

### Ultrastructure expansion microscopy (U-ExM)

Fixed ookinetes were sedimented onto 10-mm round Poly-D-Lysine coated coverslips (A3890401, Gibco) for 30 min, then prepared for U-ExM as described previously (Gambarotto et al., 2021; Zeeshan *et al*., 2022). Expanded gels were immuno-labelled using primary antibodies against α-tubulin (mouse-1:1000 dilution, Sigma) and anti-GFP (rabbit-1:200) in 3% BSA in PBS overnight. The gels were washed three times for ten minutes with PBST (Phosphate-Buffered Saline with 0.1% Tween 20) and then incubated with secondary antibodies: anti-mouse Alexa 488 (1:100 dilution, ThermoFisher) and anti-rabbit (Alexa 568 (1:1000) in PBS. The gels were washed again and expanded overnight. Images were acquired with an inverted microscope using Zeiss Plan-Apochromat 63x/1.4 Oil immersion or Zeiss C-Apochromat 63×/1.2 W Korr M27 water immersion objective on a Zeiss Elyra PS.1 microscope.

### Phenotypic analyses

To study the phenotype of Δ*PbApicortin*, approximately 60,000 parasites of either the Δ*PbApicortin* or WT-GFP lines were injected intraperitoneally (i.p.) into mice. Asexual blood stages and gametocyte production were monitored by microscopy on Giemsa-stained thin smears. Four to five days post infection, exflagellation and ookinete conversion were examined as described previously (Guttery *et al*., 2012) using a Zeiss AxioImager M2 microscope (Carl Zeiss, Inc) fitted with an AxioCam ICc1 digital camera. To analyse mosquito transmission of Δ*PbApicortin* and WT-GFP parasites, 50-60 *A. stephensi* SD 500 mosquitoes were allowed to feed for 20 min on anaesthetized, infected mice with a ∼15% asexual parasitaemia and carrying comparable numbers of gametocytes as determined on Giemsa-stained blood films. To assess mid-gut infection, approximately 15 guts were dissected from mosquitoes on day 14 post feeding and oocysts were counted on a Zeiss AxioImager M2 microscope using 10x and 63x oil immersion objectives. On day 21 post-feeding, another 20 mosquitoes were dissected, and their guts and salivary glands crushed separately in a loosely fitting homogenizer to release sporozoites, which were then quantified using a haemocytometer or used for imaging and motility assays. Mosquito bite back experiments were performed 21 days post-feeding using naive mice; 15-20 mosquitoes infected with either WT-GFP or Δ*PbApicortin* parasites were fed for at least 20 min on naive CD1 outbred mice and then infection was monitored after 3 days by examining a blood smear stained with Giemsa’s reagent. For comparison between Δ*PbApicortin* and WT-GFP, an unpaired Student’s *t*-test was used.

### Ookinete motility assays

The motility of Δ*PbApicortin* ookinetes was assessed using Matrigel as described previously (Volkmann et al., 2012; Zeeshan et al., 2020). Briefly, ookinete cultures grown for 24 h were added to an equal volume of Matrigel (Corning), mixed thoroughly, dropped onto a slide, covered with a cover slip, and sealed with nail polish. The Matrigel was then allowed to set at 20°C for 30 min. After identifying a field containing an ookinete, time-lapse videos (one frame every 5 s for 100 cycles) were collected using the differential interference contrast settings with a 63× objective lens on a Zeiss AxioImager M2 microscope fitted with an AxioCam ICc1 digital camera and analysed with the Zen lite 3.8 software (Zeiss).

### Sporozoite motility assays

Sporozoites were isolated from salivary glands of mosquitoes infected with either WT-GFP or Δ*PbApicortin* parasites on day 21 post infection. Isolated sporozoites in RPMI 1640 medium containing 3% bovine serum albumin (Fisher Scientific) were pelleted (5 min, 5,000 rpm, 4°C) and used for motility assays as described previously (Wall et al., 2019). Briefly, a drop (6 μl) of suspended sporozoites was transferred onto a microscope glass slide with a cover slip. Time-lapse videos of sporozoites (one frame every 1 s for 50 cycles) were taken using the differential interference contrast settings with a 63x objective lens on a Zeiss AxioImager M2 microscope and analysed with the Zen lite 3.8 software (Zeiss).

### Protein sequence analysis

Orthologues of Apicortin and Doublecortin were identified using InterPro (Blum et al., 2025) and aligned using Clustal Omega hosted at EMBL-EBI (Madeira et al., 2024). The structure of *P. berghei* Apicortin was predicted using the AlphaFold structural database (Jumper et al., 2021; Varadi et al., 2022).

## Acknowledgments

None

## Supplementary Materials

### Supplementary Figure

**Fig. S1. Generation and genotype analysis of transgeni*c* parasites**

**(A)** Schematic representation for 3’-tagging of *PbApicortin* gene with green fluorescent protein (GFP) sequence via single homologous recombination. **(B)** Integration PCR showing correct integration of tagging construct. **(C)** Schematic representation of the endogenous *PbApicortin* locus, the targeting gene deletion construct and the recombined locus following double homologous recombination. **(D)** Integration PCR showing correct integration with expected size of bands and deletion of *apicortin* gene from knockout (KO).

### Supplementary Table and Videos

**Table S1. Oligonucleotides used in this study**

**Video SV1: Gliding motility of WT**-**GFP ookinetes**

**Video SV2: Gliding motility** Δ***PbApicortin* ookinetes**

**Video SV3: Gliding motility of WT**-**GFP salivary gland sporozoite**

**Video SV4: Gliding motility** Δ***PbApicortin* salivary gland sporozoite**

## References

Blum, M., Andreeva, A., Florentino, L.C., Chuguransky, S.R., Grego, T., Hobbs, E., Pinto, B.L., Orr, A., Paysan-Lafosse, T., Ponamareva, I., et al. (2025). InterPro: the protein sequence classification resource in 2025. Nucleic Acids Res 53, D444–D456. 10.1093/nar/gkae1082.

Chakrabarti, M., Joshi, N., Kumari, G., Singh, P., Shoaib, R., Munjal, A., Kumar, V., Behl, A., Abid, M., Garg, S., et al. (2021). Interaction of Plasmodium falciparum apicortin with alpha- and beta-tubulin is critical for parasite growth and survival. Sci Rep 11, 4688. 10.1038/s41598-021-83513-5.

Dos Santos Pacheco, N., Tosetti, N., Koreny, L., Waller, R.F., and Soldati-Favre, D. (2020). Evolution, Composition, Assembly, and Function of the Conoid in Apicomplexa. Trends Parasitol 36, 688–704. 10.1016/j.pt.2020.05.001.

Frenal, K., Dubremetz, J.F., Lebrun, M., and Soldati-Favre, D. (2017). Gliding motility powers invasion and egress in Apicomplexa. Nat Rev Microbiol 15, 645–660. 10.1038/nrmicro.2017.86.

Fu, M.M., McAlear, T.S., Nguyen, H., Oses-Prieto, J.A., Valenzuela, A., Shi, R.D., Perrino, J.J., Huang, T.T., Burlingame, A.L., Bechstedt, S., and Barres, B.A. (2019). The Golgi Outpost Protein TPPP Nucleates Microtubules and Is Critical for Myelination. Cell 179, 132–146 e114. 10.1016/j.cell.2019.08.025.

Gambarotto, D., Hamel, V., and Guichard, P. (2021). Ultrastructure expansion microscopy (U-ExM). Methods Cell Biol 161, 57–81. 10.1016/bs.mcb.2020.05.006.

Guttery, D.S., Poulin, B., Ferguson, D.J., Szoor, B., Wickstead, B., Carroll, P.L., Ramakrishnan, C., Brady, D., Patzewitz, E.M., Straschil, U., et al. (2012). A unique protein phosphatase with kelch-like domains (PPKL) in Plasmodium modulates ookinete differentiation, motility and invasion. PLoS Pathog 8, e1002948. 10.1371/journal.ppat.1002948.

Guttery, D.S., Zeeshan, M., Ferguson, D.J.P., Holder, A.A., and Tewari, R. (2022). Division and Transmission: Malaria Parasite Development in the Mosquito. Annu Rev Microbiol 76, 113–134. 10.1146/annurev-micro-041320-010046.

Hu, K., Roos, D.S., and Murray, J.M. (2002). A novel polymer of tubulin forms the conoid of Toxoplasma gondii. J Cell Biol 156, 1039–1050. 10.1083/jcb.200112086.

Janse, C.J., Franke-Fayard, B., Mair, G.R., Ramesar, J., Thiel, C., Engelmann, S., Matuschewski, K., van Gemert, G.J., Sauerwein, R.W., and Waters, A.P. (2006). High efficiency transfection of Plasmodium berghei facilitates novel selection procedures. Mol Biochem Parasitol 145, 60–70. 10.1016/j.molbiopara.2005.09.007.

Jumper, J., Evans, R., Pritzel, A., Green, T., Figurnov, M., Ronneberger, O., Tunyasuvunakool, K., Bates, R., Zidek, A., Potapenko, A., et al. (2021). Highly accurate protein structure prediction with AlphaFold. Nature 596, 583–589. 10.1038/s41586-021-03819-2.

Koreny, L., Zeeshan, M., Barylyuk, K., Tromer, E.C., van Hooff, J.J.E., Brady, D., Ke, H., Chelaghma, S., Ferguson, D.J.P., Eme, L., et al. (2021). Molecular characterization of the conoid complex in Toxoplasma reveals its conservation in all apicomplexans, including Plasmodium species. PLoS Biol 19, e3001081. 10.1371/journal.pbio.3001081.

Legal, T., Joachimiak, E., Parra, M., Peng, W., Tam, A., Black, C., Valente-Paterno, M., Brouhard, G., Gaertig, J., Wloga, D., and Bui, K.H. (2024). Structure of the ciliary tip central pair reveals the unique role of the microtubule-seam binding protein SPEF1. bioRxiv. 10.1101/2024.12.02.626492.

Leung, J.M., Nagayasu, E., Hwang, Y.C., Liu, J., Pierce, P.G., Phan, I.Q., Prentice, R.A., Murray, J.M., and Hu, K. (2020). A doublecortin-domain protein of Toxoplasma and its orthologues bind to and modify the structure and organization of tubulin polymers. BMC Mol Cell Biol 21, 8. 10.1186/s12860-020-0249-5.

Madeira, F., Madhusoodanan, N., Lee, J., Eusebi, A., Niewielska, A., Tivey, A.R.N., Lopez, R., and Butcher, S. (2024). The EMBL-EBI Job Dispatcher sequence analysis tools framework in 2024. Nucleic Acids Res 52, W521–W525. 10.1093/nar/gkae241.

Moores, C.A., Perderiset, M., Francis, F., Chelly, J., Houdusse, A., and Milligan, R.A. (2004). Mechanism of microtubule stabilization by doublecortin. Mol Cell 14, 833–839. 10.1016/j.molcel.2004.06.009.

Nagayasu, E., Hwang, Y.C., Liu, J., Murray, J.M., and Hu, K. (2017). Loss of a doublecortin (DCX)-domain protein causes structural defects in a tubulin-based organelle of Toxoplasma gondii and impairs host-cell invasion. Mol Biol Cell 28, 411–428. 10.1091/mbc.E16-08-0587.

Olah, J., Vincze, O., Virok, D., Simon, D., Bozso, Z., Tokesi, N., Horvath, I., Hlavanda, E., Kovacs, J., Magyar, A., et al. (2011). Interactions of pathological hallmark proteins: tubulin polymerization promoting protein/p25, beta-amyloid, and alpha-synuclein. J Biol Chem 286, 34088–34100. 10.1074/jbc.M111.243907.

Orosz, F. (2009). Apicortin, a unique protein, with a putative cytoskeletal role, shared only by apicomplexan parasites and the placozoan Trichoplax adhaerens. Infect Genet Evol 9, 1275–1286. 10.1016/j.meegid.2009.09.001.

Orosz, F. (2016). Wider than Thought Phylogenetic Occurrence of Apicortin, A Characteristic Protein of Apicomplexan Parasites. J Mol Evol 82, 303–314. 10.1007/s00239-016-9749-5.

Orosz, F. (2021). Apicortin, a Constituent of Apicomplexan Conoid/Apical Complex and Its Tentative Role in Pathogen-Host Interaction. Trop Med Infect Dis 6. 10.3390/tropicalmed6030118.

Portman, N., and Slapeta, J. (2014). The flagellar contribution to the apical complex: a new tool for the eukaryotic Swiss Army knife? Trends Parasitol 30, 58–64. 10.1016/j.pt.2013.12.006.

Reiner, O., Coquelle, F.M., Peter, B., Levy, T., Kaplan, A., Sapir, T., Orr, I., Barkai, N., Eichele, G., and Bergmann, S. (2006). The evolving doublecortin (DCX) superfamily. BMC Genomics 7, 188. 10.1186/1471-2164-7-188.

Sinden, R.E., Canning, E.U., and Spain, B. (1976). Gametogenesis and fertilization in Plasmodium yoelii nigeriensis: a transmission electron microscope study. Proc R Soc Lond B Biol Sci 193, 55–76. 10.1098/rspb.1976.0031.

Sun, S.Y., Segev-Zarko, L.A., Chen, M., Pintilie, G.D., Schmid, M.F., Ludtke, S.J., Boothroyd, J.C., and Chiu, W. (2022). Cryo-ET of Toxoplasma parasites gives subnanometer insight into tubulin-based structures. Proc Natl Acad Sci U S A 119. 10.1073/pnas.2111661119.

Tewari, R., Straschil, U., Bateman, A., Bohme, U., Cherevach, I., Gong, P., Pain, A., and Billker, O. (2010). The systematic functional analysis of Plasmodium protein kinases identifies essential regulators of mosquito transmission. Cell Host Microbe 8, 377–387. 10.1016/j.chom.2010.09.006.

Varadi, M., Anyango, S., Deshpande, M., Nair, S., Natassia, C., Yordanova, G., Yuan, D., Stroe, O., Wood, G., Laydon, A., et al. (2022). AlphaFold Protein Structure Database: massively expanding the structural coverage of protein-sequence space with high-accuracy models. Nucleic Acids Res 50, D439–D444. 10.1093/nar/gkab1061.

Volkmann, K., Pfander, C., Burstroem, C., Ahras, M., Goulding, D., Rayner, J.C., Frischknecht, F., Billker, O., and Brochet, M. (2012). The alveolin IMC1h is required for normal ookinete and sporozoite motility behaviour and host colonisation in Plasmodium berghei. PLoS One 7, e41409. 10.1371/journal.pone.0041409.

Wall, R.J., Zeeshan, M., Katris, N.J., Limenitakis, R., Rea, E., Stock, J., Brady, D., Waller, R.F., Holder, A.A., and Tewari, R. (2019). Systematic analysis of Plasmodium myosins reveals differential expression, localisation, and function in invasive and proliferative parasite stages. Cell Microbiol, e13082. 10.1111/cmi.13082.

WHO (2024). World Malaria Report. https://www.who.int/teams/global-malaria-programme/reports/world-malaria-report-2024.

Yang, S., Cai, M., Huang, J., Zhang, S., Mo, X., Jiang, K., Cui, H., and Yuan, J. (2023). EB1 decoration of microtubule lattice facilitates spindle-kinetochore lateral attachment in Plasmodium male gametogenesis. Nat Commun 14, 2864. 10.1038/s41467-023-38516-3.

Zeeshan, M., Brady, D., Stanway, R.R., Moores, C.A., Holder, A.A., and Tewari, R. (2020). Plasmodium berghei Kinesin-5 Associates With the Spindle Apparatus During Cell Division and Is Important for Efficient Production of Infectious Sporozoites. Front Cell Infect Microbiol 10, 583812. 10.3389/fcimb.2020.583812.

Zeeshan, M., Rashpa, R., Ferguson, D.J.P., Abel, S., Chahine, Z., Brady, D., Vaughan, S., Moores, C.A., Le Roch, K.G., Brochet, M., et al. (2022). Genome-wide functional analysis reveals key roles for kinesins in the mammalian and mosquito stages of the malaria parasite life cycle. PLoS Biol 20, e3001704. 10.1371/journal.pbio.3001704.

Zeeshan, M., Rea, E., Abel, S., Vukusic, K., Markus, R., Brady, D., Eze, A., Rashpa, R., Balestra, A.C., Bottrill, A.R., et al. (2023). Plasmodium ARK2 and EB1 drive unconventional spindle dynamics, during chromosome segregation in sexual transmission stages. Nat Commun 14, 5652. 10.1038/s41467-023-41395-3.

