## Supplementary figures and images for "Apicortin defines the Plasmodium apical conoid body during transmission but is dispensable for the parasite life cycle"

### Fig S1

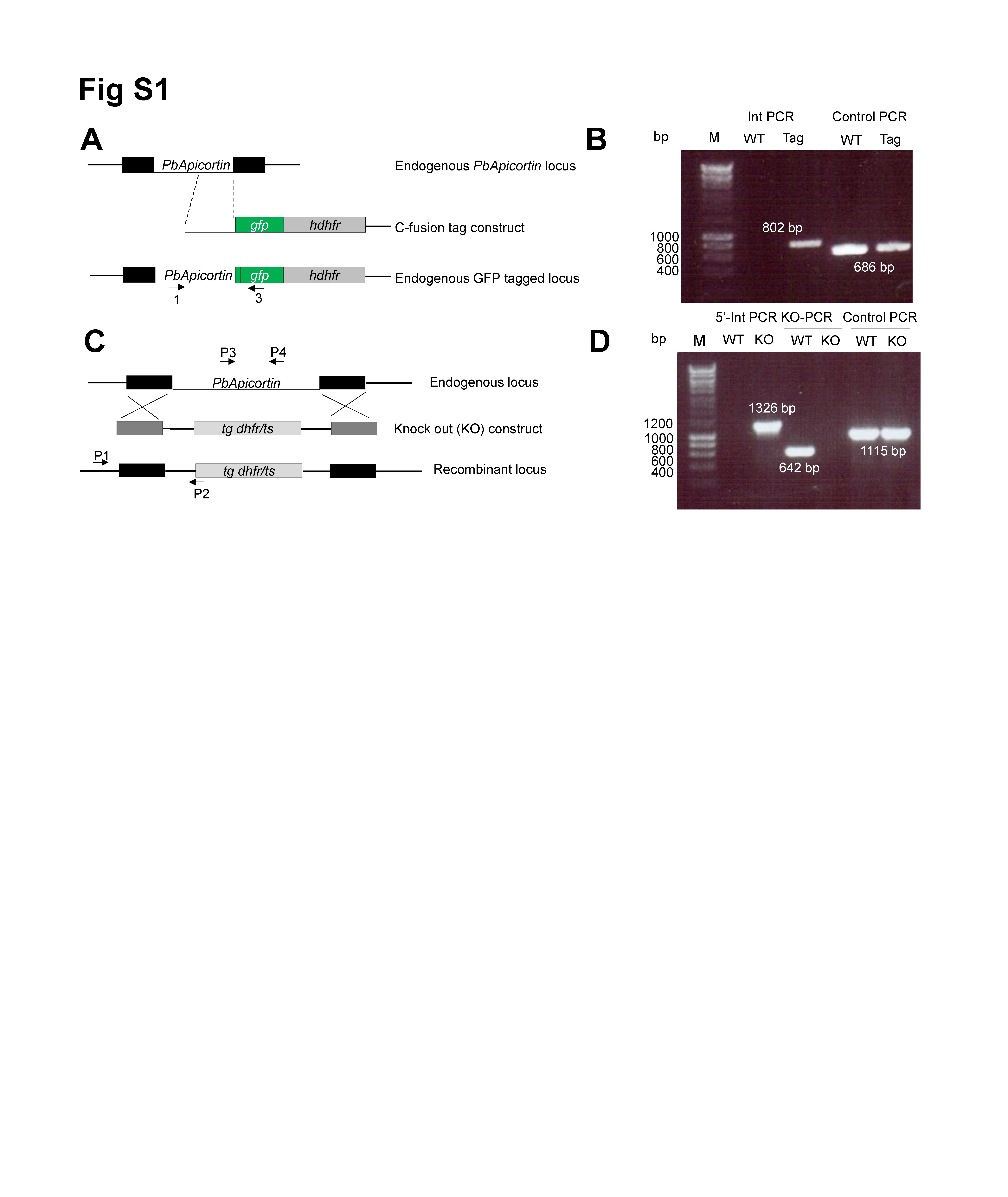
